# Whole-genome characterization in pedigreed non-human primates using Genotyping-By-Sequencing and imputation

**DOI:** 10.1101/043240

**Authors:** Ben N Bimber, Michael J Raboin, John Letaw, Kimberly Nevonen, Jennifer E Spindel, Susan R McCouch, Rita Cervera-Juanes, Eliot Spindel, Lucia Carbone, Betsy Ferguson, Amanda Vinson

## Abstract

**Background:** Rhesus macaques are widely used in biomedical research, but the application of genomic information in this species to better understand human disease is still undeveloped. Whole-genome sequence (WGS) data in pedigreed macaque colonies could provide substantial experimental power, but the collection of WGS data in large cohorts remains a formidable expense. Here, we describe a cost-effective approach that selects the most informative macaques in a pedigree for whole-genome sequencing, and imputes these dense marker data into all remaining individuals having sparse marker data, obtained using Genotyping-By-Sequencing (GBS).

**Results:** We developed GBS for the macaque genome using a single digest with *PstI*, followed by sequencing to 30X coverage. From GBS sequence data collected on all individuals in a 16-member pedigree, we characterized an optimal 22,455 sparse markers spaced ~125 kb apart. To characterize dense markers for imputation, we performed WGS at 30X coverage on 9 of the 16 individuals, yielding ~10.2 million high-confidence variants. Using the approach of “Genotype Imputation Given Inheritance” (GIGI), we imputed alleles at an optimized dense set of 4,920 variants on chromosome 19, using 490 sparse markers from GBS. We assessed changes in accuracy of imputed alleles, 1) across 3 different strategies for selecting individuals for WGS, i.e., a) using “GIGI-Pick” to select informative individuals, b) sequencing the most recent generation, or c) sequencing founders only; and 2) when using from 1-9 WGS individuals for imputation. We found that accuracy of imputed alleles was highest using the GIGI-Pick selection strategy (median 92%), and improved very little when using >4 individuals with WGS for imputation. We used this ratio of 4 WGS to 12 GBS individuals to impute an expanded set of ~14.4 million variants across all 20 macaque autosomes, achieving ~85-88% accuracy per chromosome.

**Conclusions:** We conclude that an optimal tradeoff exists at the ratio of 1 individual selected for WGS using the GIGI-Pick algorithm, per 3-5 relatives selected for GBS, a cost savings of ~67-83% over WGS of all individuals. This approach makes feasible the collection of accurate, dense genome-wide sequence data in large pedigreed macaque cohorts without the need for expensive WGS data on all individuals.

## BACKGROUND

The analysis of whole-genome sequence data in non-human primates (NHPs) can play a significant role in advancing the application of genomic medicine to human disease. Potential uses of these data include the identification of novel genetic variants that influence conserved pathways of disease pathology, the development of novel therapeutics that target these variants, and the characterization of variants that influence efficacy and response to therapeutics. Given their high degree of genetic and physiological similarity to humans, and their ubiquity in biomedical research, it is surprising that the use of the rhesus macaque for these purposes has been so slow to develop. One likely reason for this delay is the dearth of genome-wide sequence information on sufficient numbers of animals to support such studies, which typically require large numbers of phenotyped and genotyped subjects. However, the collection of dense sequence data in large cohorts remains a formidable expense, and a cost-effective solution to this problem is needed if we are to reap the full benefit of non-human primate (NHP) models in both basic science and preclinical research.

Whole-genome sequencing (WGS) of large cohorts remains a very expensive undertaking, both now and likely long after we achieve the $1,000 per genome benchmark. Several sequencing strategies have been developed to address this problem, each of which strikes a different balance between sequencing costs, sequence depth, and coverage across the genome. Although deep WGS is the most unbiased and comprehensive method for surveying genetic variants [1], at approximately $3000/genome for standard 30X coverage, its cost remains prohibitive in the foreseeable future for large cohort studies. A second strategy aims to cover the whole genome but at greatly reduced depth (i.e., “low coverage sequencing”), which lowers costs to $200-600/genome. However, this approach reduces the accuracy of resulting genotype data, particularly for smaller studies of rare or low-frequency variants [2]. Another strategy is to sequence only a portion of the genome, i.e., “reduced representation” approaches, which offers a reasonable compromise between sequencing depth and breadth of coverage. The most common of the reduced representation approaches is whole-exome sequencing, currently ~$300/genome for 30X coverage, in which a commercial hybridization kit is used to capture genomic fragments enriched for exons in protein-coding genes. While this approach produces coverage of genomic regions that are of interest to many Mendelian diseases, coverage of regulatory elements or other non-coding regions is sacrificed. Moreover, most commercial exome capture tools are designed for humans or rodents, and thus will miss some portion of the NHP exome.

More recently, a reduced representation approach called genotyping-by-sequencing (GBS) has lowered the cost per genome dramatically, by taking advantage of classical molecular biology methods that capture a more evenly distributed subset of the genome. In the GBS method, standard restriction enzymes target conserved cut sites that span the genome, and the resulting fragments are sequenced to the desired coverage. While these fragments still represent only a small portion of the genome, they can be distributed more evenly than in other methods such as exome capture. Importantly, because the GBS approach does not require proprietary capture technology and can be highly multiplexed, costs can be reduced to as little as $50 per genome and this approach can be applied to species that lack available commercial arrays. This approach has been applied to many agricultural and other economically important species to construct dense genetic linkage maps and identify QTLs [3-8], to improve genome assemblies [9], and to investigate population structure, diversity, and evolutionary history [10-12].

Further gains in the amount of sequence information obtained at the lowest possible cost could be achieved by combining GBS data with imputation, particularly for NHP cohorts with pedigree information. In this approach, whole-genome sequence data collected in selected individuals within the pedigree are used to impute dense genotypes into their many relatives, in which only sparse genotype data (e.g., obtained by GBS) has been collected. These sparse data from GBS are used to anchor the imputation of genotypes at intervening and more densely spaced loci across the genome, by leveraging information on expected allele-sharing among relatives. This strategy is appealing for many captive NHP breeding colonies, where deep and well-defined pedigrees could permit extremely cost-effective, whole-genome characterization. However, the selection of the most informative animals in the pedigree for whole-genome sequencing is expected to have a large impact on the success of this approach, and studies addressing optimal selection strategies have only been published for human pedigrees, which are typically much smaller and less complex than those characterized for NHP cohorts.

Although whole-genome sequencing combined with GBS and imputation presents a significant opportunity for obtaining dense sequence data at minimal cost, this approach has not yet been applied to a pedigreed NHP cohort. Thus, our objectives were to, 1) develop a reliable GBS method in the macaque genome to support pedigree-based imputation; 2) to assess the extent and accuracy of imputed dense marker data from WGS using sparse marker data from GBS; and 3) to compare the accuracy of imputed dense marker data among different strategies commonly used to select individuals for WGS, and among different ratios of WGS to GBS individuals in the pedigree. Here, we show that a *PstI* digest in the macaque genome produces >22,000 high-quality sparse variants that are suitable for use in imputation. We further show that the pedigree-based “Genotype Imputation Given Inheritance” imputation approach (i.e., “GIGI”; [13]), combined with the GIGI-Pick method [14] of selecting individuals for WGS, allowed us to impute >14 million variants throughout a 16-member pedigree with ~85-88% accuracy, using only 4 individuals with WGS and 12 individuals with GBS. This strategy represents a reasonable tradeoff between sequencing costs, and the amount and quality of dense sequence data obtained on as many individuals as possible.

## METHODS

### Animal care and welfare

All macaque samples used in this study were collected during routine veterinary care procedures approved by the Institutional Animal Care and Use Committee of the Oregon Health & Science University (Protocol Number: IS00002621); these samples are part of the much larger ONPRC DNA Biobank. Animal care personnel and staff veterinarians of the ONPRC provide routine and emergency health care to all animals in accordance with the Guide for the Care and Use of Laboratory Animals, and the ONPRC is certified by the Association for Assessment and Accreditation of Laboratory Animal Care International.

### Pedigree configuration and validation

We selected 16 closely related animals from the larger Oregon National Primate Research Center (ONPRC) colony pedigree as the focus of this study (see Fig. 1). These animals were selected to represent the most common relationships in the colony, including parent/offspring, half-sibling, half-avuncular, half-cousin, and grandparent/grandchild relationships. Because assumed pedigree relationships may prove to be incorrect when comprehensive genotype data are examined, we explored the accuracy of our focal 16-member pedigree using a set of ~5,000 markers on chromosome 19 generated from our GBS sequencing experiments (see *Imputation accuracy on chromosome 19 across selection strategies*, below), employing algorithms that assess Mendelian consistent error both pairwise between relatives and within families, as implemented in PedCheck [15] and GIGI-Check [16] software. No significant departures from expected patterns of allele-sharing were noted, confirming the validity of the pedigree configuration depicted in Fig. 1. Nine animals were selected using an *ad hoc* approach for whole-genome sequencing in this study, based on their position within the pedigree.

**Figure 1.**
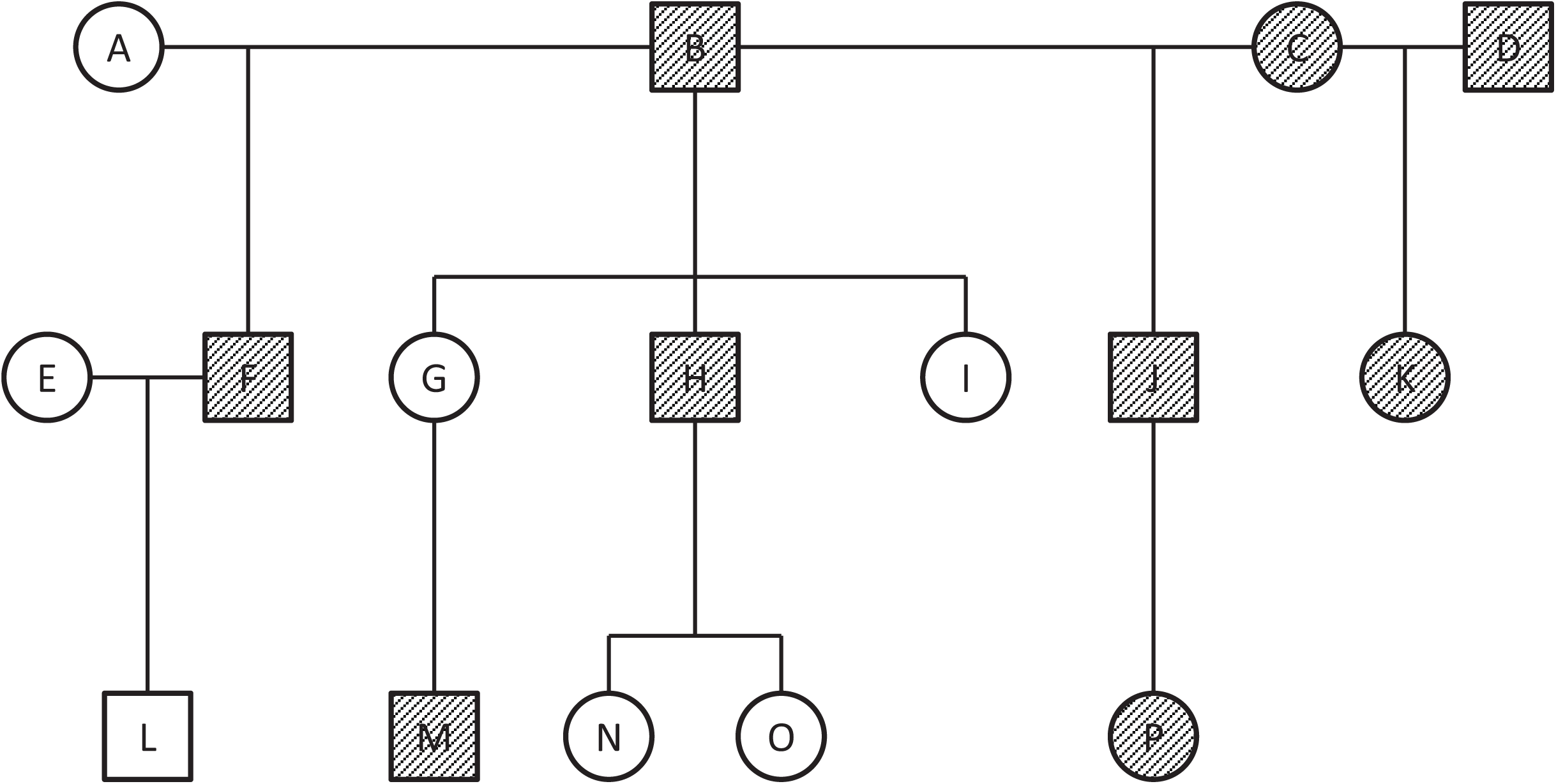
Pedigree diagram. Pedigree diagram of the 16 subjects included in this study. Subjects with whole genome sequence data are shown in gray; all subjects have GBS data.

### Genomic DNA Isolation and Quantification

Genomic DNA (gDNA) was extracted from 3 ml of whole blood using the ArchivePure DNA Blood Kit (5 Prime, Inc.), following the manufacturer’s recommendations. Genomic DNA was quantified with the Qubit High Sensitivity dsDNA Assay (Life Technologies, CA).

### Genotyping-By-Sequencing (GBS)

To determine the optimal restriction enzymes for conducting GBS in rhesus macaques, we first performed *in silico* prediction of cut sites using the most recent build of the macaque genome [17] that would be expected to produce 60,000-100,000 DNA fragments in the 200-500 bp size range, while also minimizing the presence of repeat sequences (e.g. retrotransposons, DNA satellites). We initially tested the enzymes *ApeKI, BglII, EcoRI, HindIII, PspXI, PstI*, and *SalI*, ultimately selecting *BglII* and *PstI* as the two enzymes most likely to meet these criteria. We then generated GBS libraries based on these 2 enzymes using a modified version of the method described by Elshire et al. [18]. Specifically, to create the adaptors, oligonucleotides for the top and bottom strands for each barcoded adaptor and for the two common adaptors (one for *BglII* and one for *PstI)* were paired and annealed in 1X Annealing Buffer (20mM NaCl, 10mM Tris-HCl pH 7.5, 2mM MgCl_2_) using a thermal cycler (3 min at 95C, ramp down 1.6C/min for 44 cycles, cool to 4C). All adaptors were quantified with the Qubit Broad Range dsDNA Assay (Invitrogen). Each of the 32 barcoded adaptors was then paired with a common adaptor at a 1:1 ratio. Each of the 16 genomic DNAs was digested with *BglII* and *PstI* in separate reactions. All 32 reactions (500ng DNA, 10U enzyme, in 20uL volume) were incubated for 2 hours at 37°C, and digests were ligated (400U T4 DNA Ligase (New England Biolabs) to adaptor mixes (4.5ng *BglII*, 15ng *PstI*, in 50ul volume) for 1 hour at 22°C. Four (4) ul from each ligation reaction was combined into two separate pools, one per enzyme. Both pools were cleaned with DNA Clean and Concentrator (Zymo Research) and eluted in 50uL. Following amplification parameters in Elshire et al [18], PCR was performed on 10 ul of each pool (Q5 High Fidelity 2X MM (New England Biolabs), 25 pmol of each primer, in 50 ul volume) using Primers A and B, to extend and complete the sequencing adaptors.

Libraries were purified using the Qiaquick PCR Purification Kit (Qiagen), quantified with the Qubit High Sensitivity dsDNA Assay (Invitrogen) and validated with the Bioanalyzer High Sensitivity Assay (Agilent). A one-sided 0.8X size selection with AMPure XP beads (Beckman-Coulter) was used to enrich larger size fragments. Libraries were sequenced on an Illumina NextSeq at the Oregon Health & Science University Integrated Genomics Laboratory to produce 30X coverage.

### Whole Genome Sequencing

Per sample, 1 μg of gDNA was sheared using a Bioruptor UCD200 (Diagenode, Denville, NJ), generating fragments around 300 bp. Libraries were constructed using the NEXTflex DNA Sequencing Kit and NEXTflex DNA barcodes (BIOO Scientific, Austin,TX) following the manufacturer’s instructions. Briefly, the ends of the sheared gDNA were repaired and adenylated, then ligated to barcoded adaptors using the reagents provided. Next, fragments of 200-400 bp were excised from a 1% agarose gel. The products were amplified by PCR using 8 cycles, then purified using 1X AMPure XP beads (Beckman-Coulter). The final libraries were quantified with the Qubit High Sensitivity dsDNA Assay (Life Technologies, CA) and validated using the 2100 Bioanalyzer High Sensitivity Assay (Agilent Technologies, Santa Clara, CA). Libraries were sequenced on a HiSeq3000 at the Oregon State University Center for Genome Research and Biocomputing, to produce 30x coverage with paired-end, 150 bp reads.

### Analysis of Sequence Data

Both whole genome and GBS data were processed using the best practice recommendations from the Broad Institute’s Genome Analysis Toolkit (GATK; [19, 20]), adapted for rhesus macaque. Briefly, paired-end reads were trimmed using Trimmomatic [21], and aligned to the most current rhesus macaque reference genome, “MacaM” [17], using Burrows-Wheeler Aligner [22]. GATK’s HaplotypeCaller was used to produce VCF files, followed by genotype calling using GenotypeGVCFs. The resulting VCF was filtered for quality, strand bias, and proximity to the read end. Additional hard filters include removal of, 1) singlenucleotide variant (SNV) clusters within a 20 bp span, 2) SNVs located within regions with greater than twice the mean coverage (potential CNV or mismapped reads), 3) SNVs that display non-Mendelian inheritance, and 4) SNVs located within repetitive regions. The analyses also employed Picard tools [23] and FASTQC [24] for quality control of the raw data, JBrowse [25] to visualize data, and BEDTools [26] to evaluate SNV and imputation marker distribution. Sequence data were managed and analyzed using DISCVR-Seq [27], a LabKey Server-based system [28].

### Sequencing and imputation strategy

We focused on chromosome 19 as a test case in order to develop an analytical pipeline that could be applied to all the remaining chromosomes. We performed imputation using the method of GIGI (“Genotype Imputation Given Inheritance”; [13]), as this method has been successfully used to impute genotypes with high accuracy in extended human pedigrees. This approach infers inheritance vectors (IVs, representing shared chromosomal segments) at sparse marker locations conditioned on observed sparse marker genotypes, and then infers IVs at dense marker locations conditioned on the sparse marker IVs, together with the genetic map. The probability distribution is then estimated for each missing genotype at a dense marker position, conditioned on observed genotypes at all dense marker positions, corresponding allele frequencies, and IVs corresponding to dense markers. In the last step, genotypes may be called using these estimated probabilities, based on user-defined thresholds. We estimated inheritance vectors according to the algorithm of [29], as implemented using a Markov-Chain Monte-Carlo (MCMC) sampler in the gl_auto function in the software package for genetic epidemiology MORGAN 3; available at[30]. The GIGI approach has been implemented in a software package of the same name, and is available at [31].

To characterize the sparse set of markers needed to facilitate imputation of dense marker genotypes on chromosome 19, we identified a set of markers that could be detected reliably by GBS for as many macaques as possible. Accordingly, we selected a set of high-quality SNVs that, 1) were sequenced to at least 20X depth across the majority of GBS libraries, 2) were spaced evenly across the genome, 3) had minor allele frequencies (MAF) >0.25, and 4) were in excess of what was needed to meet the desired goal of ~0.5-1.0 cM average marker spacing. We refer to these as “framework” markers, as discussed in Cheung et al., 2013 [13]. Using this approach, the desired spacing can be maintained in an approximate fashion, even when individuals are missing a substantial amount of genotype data, an outcome characteristic of the GBS method [32]. Second, we selected a non-overlapping set of SNVs from our WGS data that were more densely distributed than the framework markers, designated as our “dense” markers, and which we attempted to impute into animals having only sparse framework marker genotypes from GBS. These dense markers were selected from the set of all high-confidence SNVs identified in our cohort.

To determine the success of imputing dense marker data into animals having only sparse framework marker data, we evaluated the accuracy of imputed alleles for each recipient based on their framework marker data obtained by GBS. Here, we define accuracy as the proportion of alleles imputed correctly among all attempted allele calls at that position, such that a correctly imputed allele is concordant with the allele call from either WGS or GBS sequence data. We additionally define rare variants as those having only 1 copy among a total of 30 chromosomes (i.e., singletons, present at ~3% frequency) with WGS data that we have studied to date, including the 9 individuals discussed in this paper and an additional 6 unrelated rhesus macaques (unpublished data). We define accuracy of imputation for rare variants as the proportion of rare alleles imputed correctly among all rare variant heterozygotes called from either WGS or GBS sequence data.

We assessed accuracy of imputed variants over a range of 1-9 animals with dense marker data from WGS, imputed into the remaining pedigree members using only sparse data from GBS. Individuals with WGS data were added consecutively in the following order: B, H, J, F, M, K, P, C, D (SEE FIG. 1). Thus, the first scenario used only the most informative animal (B) with WGS to impute genotypes into the remaining 15 animals within the pedigree. Subsequent scenarios retained the previous animal(s), added the next most informative animal, and imputed genotypes into the remaining animals within the pedigree. This procedure was conducted iteratively, until all 9 animals with WGS were used to impute genotypes into the remaining 7 animals in the pedigree. We used the GIGI-Pick algorithm [14] to rank our 9 animals with WGS. This algorithm calculates a metric of coverage, defined as the expected percentage of allele copies called for a variant at a random locus, conditional on fixed IVs for a specific choice of individual(s), and then iteratively selects those individuals with the highest coverage, calculated by integrating over all possible genotype configurations within a given pedigree. This algorithm is implemented in the suite of software based on the GIGI approach, and is available at [33]. We evaluated accuracy of imputed genotypes for each of our recipient macaques by masking all non-framework genotypes in recipient animals, and comparing imputed genotypes to masked genotypes obtained from either WGS or GBS data, depending on the data available for each recipient. Specifically, imputed genotypes were compared to genotypes from WGS where available, but for recipients with only GBS data available, imputed genotypes were compared to genotypes at any SNVs with coverage by GBS that were not designated as framework markers. Imputed genotypes were called as the most probable genotype at that position, using allele frequencies established from all Indian-origin rhesus macaques sequenced to date at the ONPRC as a reference (n=15, including the 9 animals from this study and 6 additional unrelated animals).

To evaluate differences in imputation success associated with different sequencing strategies, we compared the accuracy of genotypes imputed by GIGI on chromosome 19, among 3 different sequencing strategies, including GIGI-Pick and two common heuristic methods. These 2 heuristic methods include whole-genome sequencing of, 1) pedigree founders only, or 2) the most recent generation (i.e., individuals typically located at the bottom of the pedigree). To compare the different selection strategies, we examined accuracy for the scenario in which dense markers from 3 animals selected for WGS are imputed into the remaining 13 pedigree members with GBS data, based on using individuals B, H, and J (GIGI-Pick selections), B, C, and D (“Founders”), and M, P, and K (“Pedigree bottom”) strategies (see Fig. 1). Imputed genotypes were called using 2 different strategies available in the GIGI algorithm, i.e., genotypes were either above the default probability threshold, or simply as the most probable genotype at that position (“Threshold” vs. “Most Likely”, respectively), using allele frequencies established from all Indian-origin rhesus macaques sequenced to date at the ONPRC as a reference. To assess the utility of the GIGI imputation approach for imputing low-frequency variants, we further evaluated genotype accuracy within the GIGI-Pick strategy of 3 WGS into 13 GBS described above.

## RESULTS

### Whole-genome Sequencing and Variant Calling

We obtained an average 566,035,688 read pairs per sample (range 495,617,772-735,313,000) for each of the 9 individuals with WGS. These reads were aligned to MacaM, the most recent macaque genome build [17], to produce an average 27X coverage across the genome (range 24-33X). From these reads, a total of 10,193,425 high-confidence SNVs were identified across all 9 individuals, with an average of 5,037,341 variants detected per individual. The transition/transversion ratio (Ti/Tv) observed in this study was 2.17, consistent with observations in larger macaque cohorts (unpublished data). This set of sites served as the source of our optimal dense marker set, as described in Methods.

### Genotyping-By-Sequencing and Variant Calling

For each of the 16 pedigree members, we prepared and sequenced GBS libraries based on individual digests for both *BglII* and *PstI*. Among *BglII* libraries, we obtained an average 3,754,352 reads per sample, resulting in an average 4,851,004 base-pairs (bp) from 42,919 fragments with at least 20X coverage per sample (equivalent to 0.17% of the genome). In contrast, among *PstI* libraries, we obtained an average 5,686,709 reads per sample, resulting in an average 10,682,162 bp from 130,247 fragments with >20X coverage per sample (equivalent to 0.38% of the genome) (Fig. 2A-2B). Notably, although the *PstI* libraries originally had ~1.5X more reads than *BglII* libraries, they had >3-fold the number of fragments with high-depth coverage. Virtually all GBS fragments were adjacent to their predicted restriction sites, but a small number appeared to be distant from these sites (Fig. 2C-D). While these results may reflect off-target sequencing, it is also possible that they reflect restriction sites in the genome of one or more individuals not predicted by the current macaque reference genome.

**Figure 2.**
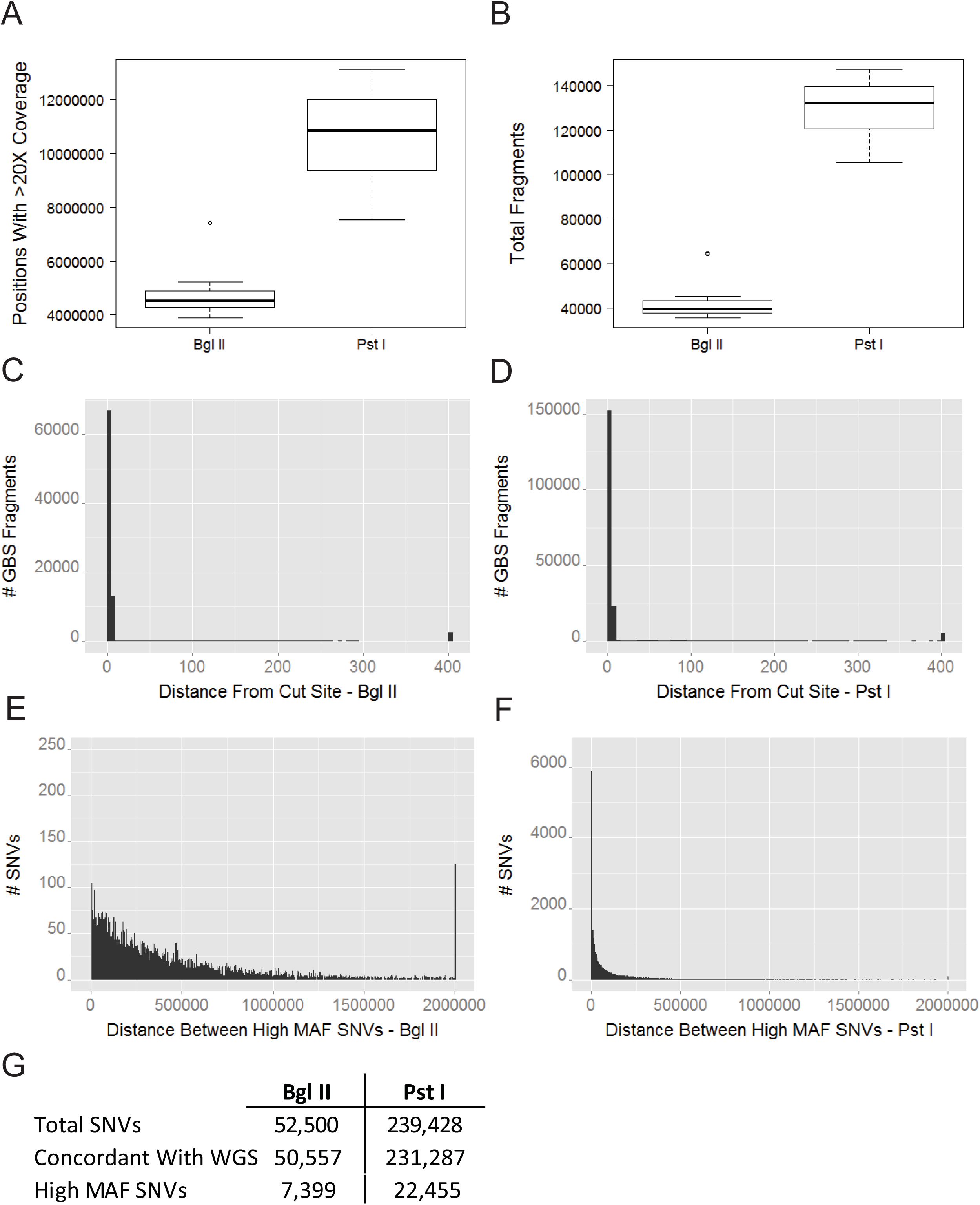
Evaluation of GBS library coverage and SNVs. **a)** The number of positions with at least 20X coverage; **b)** Total contiguous fragments with >20X coverage; **c)** Distance between each GBS fragment and nearest predicted cut site for the *BglII* libraries (all fragments >400 bp are grouped into a single bin); **d)** Distance between each GBS fragment and the nearest predicted cut site for *PstI* libraries; **e)** Distance between high MAF (>0.25) SNVs in *BglII*; **f)** Distance between high MAF SNVs in *PstI*; **g)** Total SNVs detected per enzyme, total that were concordant with WGS data, and total SNVs with MAF >0.25.

We next determined the number of high-quality SNVs available from each digest that would be suitable for use as framework markers in imputation. We first restricted SNVs to those with >20X coverage, leaving a total of 52,500 high-confidence variants in the *BglII* samples, and 239,428 variants in *PstI* samples. We further restricted these SNVs to only those that were concordant between WGS and GBS data, which included 96.3% of SNVs for *BglII* (range 95.9-97.2% among individuals) and 96.6% of SNVs for *PstI* (range 96.1-97.2% among individuals). To maximize the probability of the variant being present in as many animals as possible, we also restricted SNVs to only those with MAF >0.25, which further reduced these numbers to 7,399 variants for *BglII* and 22,455 variants for *PstI*, with an average distance between SNVs of 376,818 bp and 125,280 bp, respectively (see Fig. 2E-F). We were not able to call genotypes in all individuals for all SNV sites, due to variation among individuals in sequence quality at each site. From the *PstI* data, all individuals had sufficient data to call genotypes at an average 15,516 (~69% of 22,455) of these SNVs, but only an average of 4,418 (~60% of 7,399) of these sites could be called for all individuals from the *BglII* data. Based on the significantly greater numbers of high-quality SNVs across the macaque genome available from *PstI* sequence data, we chose this enzyme for all final imputation analyses.

### Imputation accuracy on chromosome 19 across 3 different selection strategies

Our initial tests of imputation focused on chromosome 19. To characterize the set of framework markers needed to facilitate imputation on this chromosome, we selected 490 variants spaced ~100 kb apart, from the set of 981 high-MAF variants on this chromosome, as described above (Fig. 2G). To characterize the set of dense markers to be imputed on this chromosome, we selected 4,920 variants spaced ~10 kb apart and which were not framework markers, from the set of 260,000 SNVs located on this chromosome (see above). We additionally removed a set of 578 markers that consistently performed poorly in imputation. This reduced and optimized set of dense markers on chromosome 19 was used in all imputation analyses on this chromosome, in order to limit the computational time required.

Using these chromosome-specific framework and dense marker sets, we evaluated the “Bottom of Pedigree”, the “Founders”, and the “GIGI-Pick” strategies for selecting the 3 most informative of the 9 individuals with WGS data, followed by imputation of dense markers into the remaining 13 individuals, based on their framework marker data from GBS. For both genotype-calling methods, the GIGI-Pick selection strategy produced slightly higher median accuracy of imputed genotypes than either of the other strategies, at 89.5% (“Most Likely” method, ML) or 90.1% accuracy (“Threshold” method, THR), compared to median accuracy of 88.4% (ML) or 88.8% (THR) in the “Bottom of Pedigree” strategy, and 88.6% (ML) or 87.6% (THR) in the “Founders” strategy (Fig. 3A-B). However, more individuals in the “GIGI-Pick” strategy displayed greater genotype accuracy than in either of the other 2 strategies (interquartile range (IQR) from 89.5%-92.6% for GIGI-Pick, compared to 84.2-90.1% for “Bottom of Pedigree”, and 85.9-89.5% for “Founders”). While the difference in median accuracy estimated by both genotype calling methods was <1%, 100% of genotypes were called in the ML method, while only 47.7% were called using the THR method across all strategies. Thus, the ML method called genotypes at an average 2,350 more markers per subject, while maintaining nearly identical accuracy. The GIGI-Pick strategy did produce one individual that consistently displayed much lower genotype accuracy than all other individuals; individual D had only 72.9-75.6% of genotypes accurately imputed, depending on calling method. Neither the “Bottom of Pedigree” nor the “Founders” strategy produced any individuals with substantially lower genotype accuracy than other pedigree members. Finally, we used the GIGI-Pick results to examine accuracy of imputed singleton alleles, as defined in Methods. These rare alleles were imputed a total of 405 times, representing 178 distinct alleles, with an accuracy of 93%.

**Figure 3.**
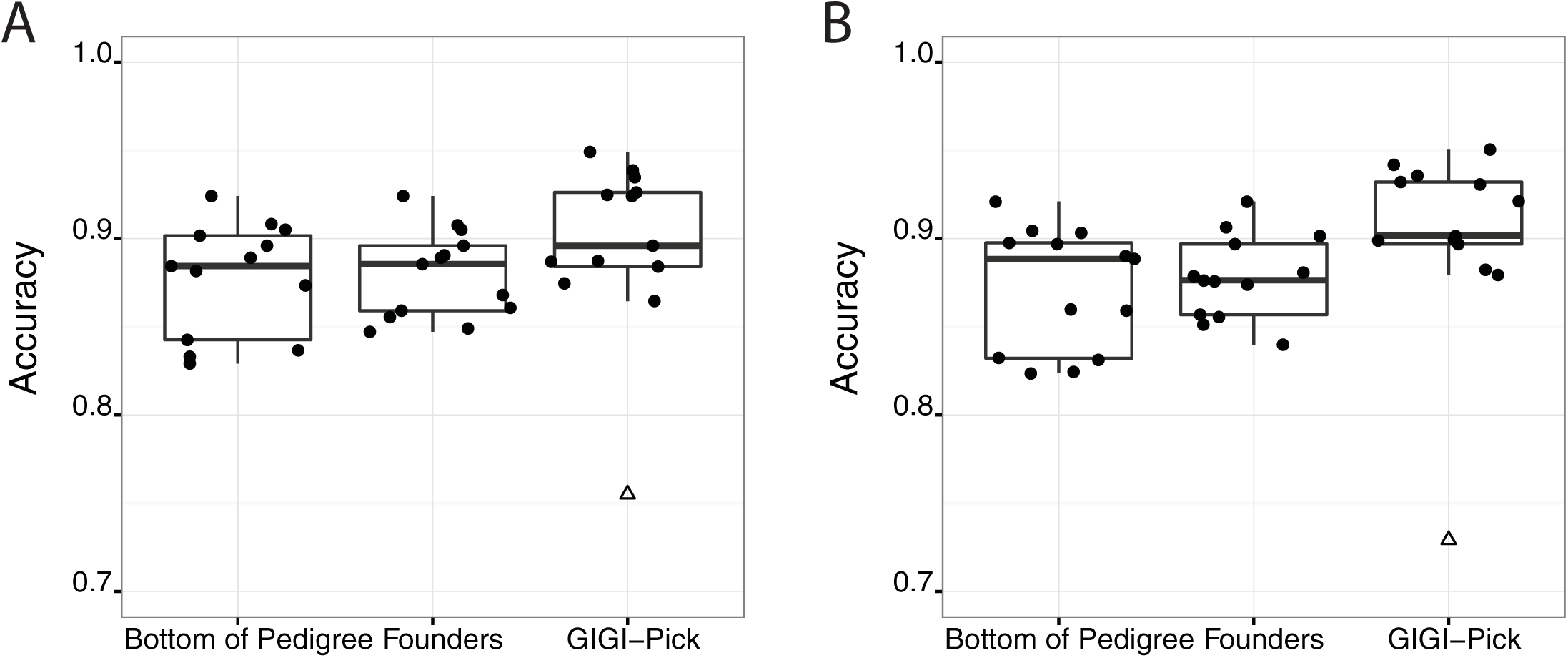
Imputation accuracy on chromosome 19, among different strategies for selecting 3 individuals for WGS. Comparison of imputation accuracy among 3 different strategies for selecting 3 individuals for WGS within the 16-member pedigree: “Bottom of Pedigree” (subjects M, P, K), “Founders” (subjects B, C, D), and “GIGI-Pick” (B, H, J). Imputation of an optimal set of dense markers was conducted for chromosome 19 from the 3 individuals with WGS, into the 13 recipient individuals with GBS data, using the GIGI imputation algorithm with the “Most-Likely” (A) and “Threshold” (B) genotype calling methods. Circles indicate accuracy for each of the 13 individuals; triangles indicate outlier individuals.

### Imputation accuracy on chromosome 19 across 9 sequencing ratios

Using the same chromosome 19 framework and dense marker sets, we used the GIGI algorithm with the ML genotype-calling method to impute our dense markers from 1-9 individuals with WGS into the remaining 7-15 pedigree members, as described in Methods. Among all imputation scenarios that added consecutively from 1 to 9 individuals with WGS, median accuracy among recipients increased from 86.3% to 93.2%, while variation in accuracy decreased (IQR from 5% at N=1 WGS, to 1% at N=9 WGS). Variability in accuracy improved in a stepwise fashion; greater variability tended to occur in parallel with an increase in median accuracy, but would improve during the following scenario in which the greater accuracy was retained or further increased. Individual D was consistently an outlier across 8 out of 9 scenarios, from 1 through 5 at 76% accuracy, then increasing to 84% accuracy in scenarios 6-8, due to the inclusion of WGS data from K, the child of D. Median accuracy surpassed 90% beginning at N=4 individuals with WGS; although variation in accuracy continued to decrease across all remaining scenarios, only slight gains in median accuracy were observed beyond this sequencing ratio (i.e., imputing from 4 individuals with WGS into 12 individuals with GBS) (Fig. 4).

**Figure 4.**
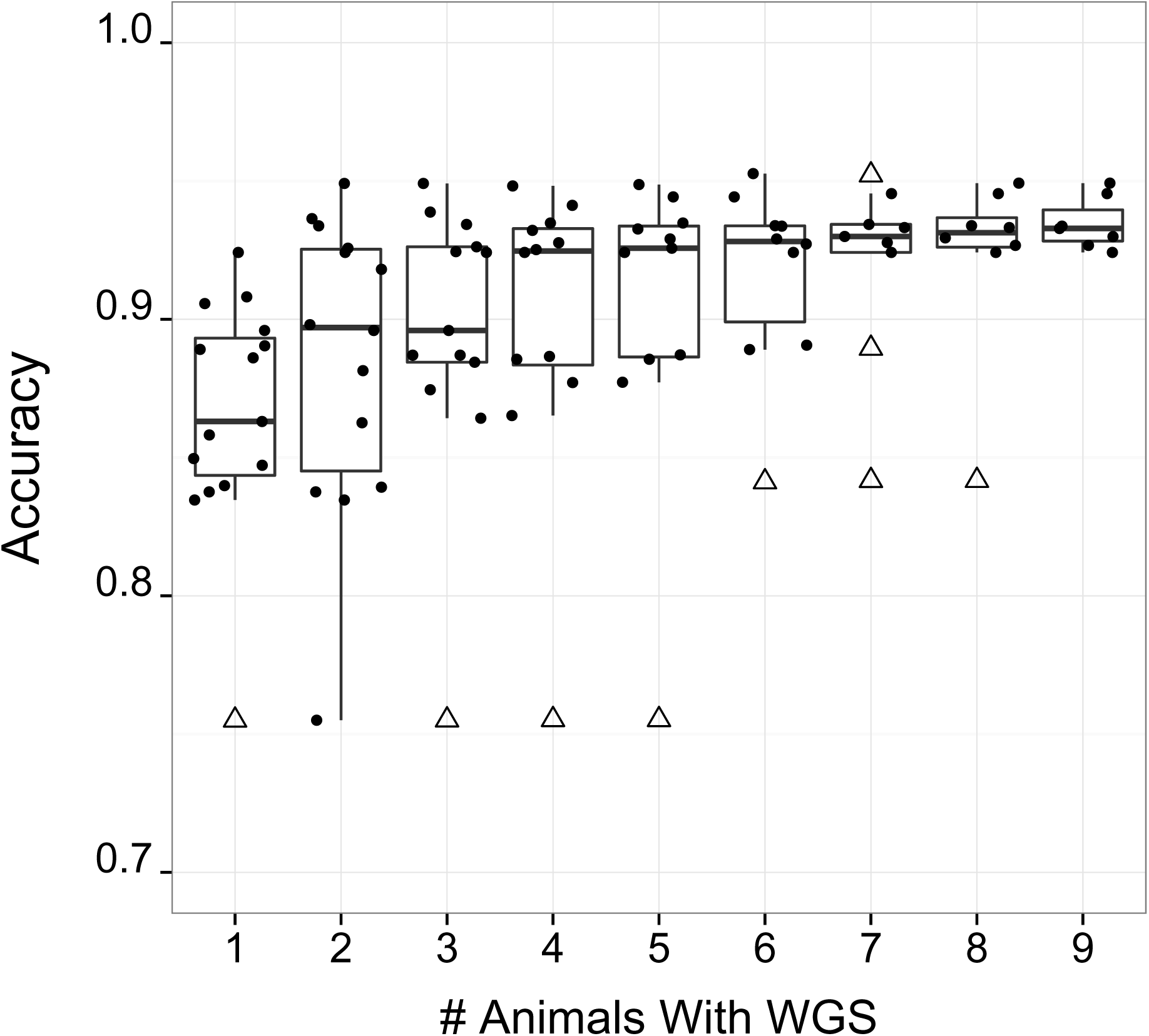
Imputation accuracy on chromosome 19 by total number of individuals selected for WGS. Accuracy for an optimal set of dense markers on chromosome 19, using 1-9 individuals with WGS data, imputed into all remaining pedigree members with GBS data, using the “most likely” genotype calling method in the GIGI algorithm [13]. Circles indicate accuracy for each individual; triangles indicate outliers. Individuals with WGS data were selected by the GIGI-Pick algorithm [14] and used for imputation in the following order: B, H, J, F, M, K, P, C, D (see Fig. 1).

### Imputation of dense markers across the genome

To evaluate our imputation strategy across the whole genome, we employed the same criteria outlined above to generate framework marker sets for each of the 20 macaque autosomes. The number of framework markers per chromosome ranged from 350 to 636, with mean spacing between framework markers among all chromosomes of ~273 kb (108 kb-467 kb). At this stage, we expanded our dense marker set to a total 14,384,988 SNVs across the macaque genome, by including SNVs discovered previously from whole-genome sequence data in an additional 6 unrelated ONPRC animals (in prep). Using the strategy identified by our analyses as the one most likely to maximize genotype accuracy while minimizing overall costs, we used the first 4 individuals among our 9 with WGS ranked by the GIGI-Pick algorithm, and imputed our expanded dense marker data into the remaining 12 pedigree members, based on the ML call method (Fig 5). Per chromosome, median accuracy ranged from 85-88%; this accuracy is somewhat lower than the ~92% accuracy achieved in our original experiment using chromosome 19. However, unlike our previous analysis that imputed a much smaller set of dense markers on chromosome 19, our final genome-wide imputation included all known variants. As before, D had consistently lower genotype accuracy at 73-77%. Individual E was also an outlier on multiple chromosomes, with accuracy ranging from 79-84%.

**Figure 5.**
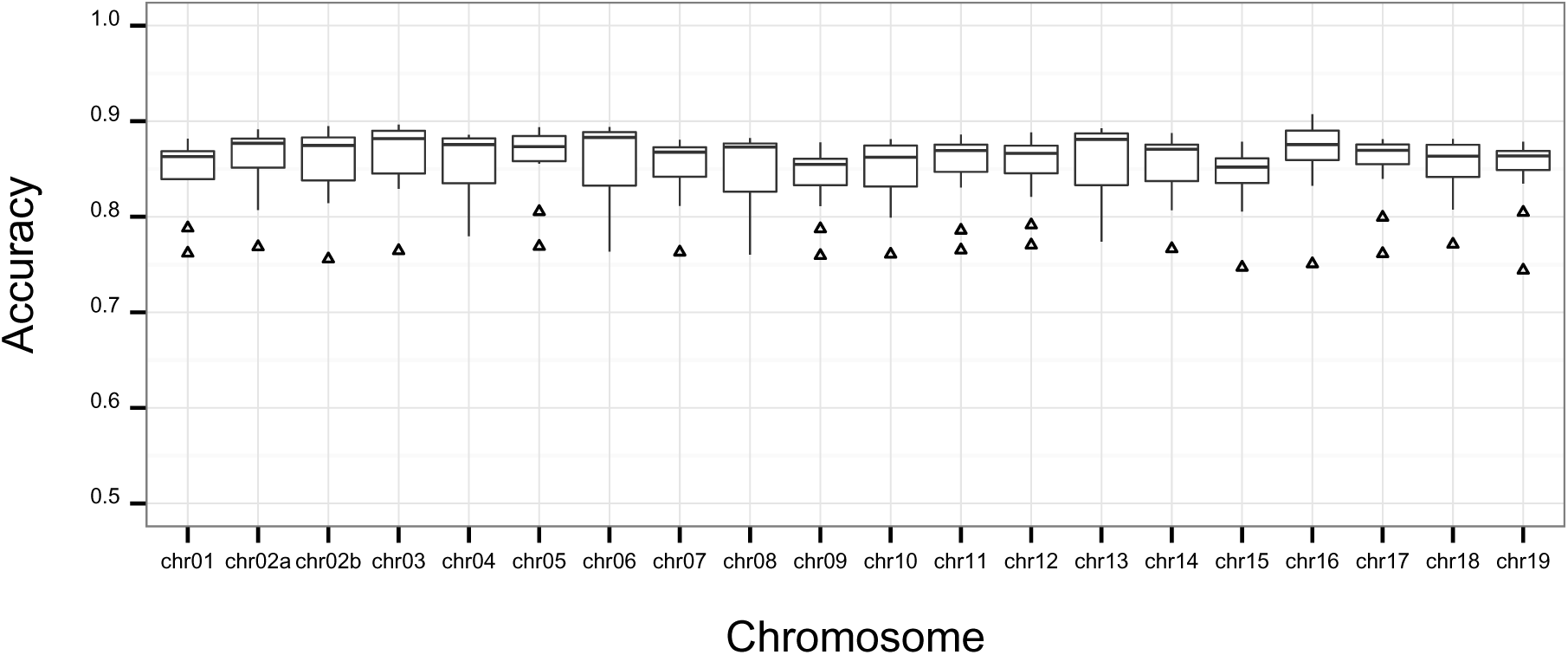
Imputation accuracy across the genome at an expanded set of dense markers. Accuracy of alleles imputed across all autosomes at 14,384,988 dense markers. Data represent accuracy of alleles imputed at dense markers for 12 pedigree members with GBS data, imputed from individuals B, H, J, and F. Alleles were called using the “most likely” genotype calling method in the GIGI algorithm [13]. Triangles indicate outliers.

## DISCUSSION

The rhesus macaque is widely used in academic biomedical research, primarily due to its utility as a model of human HIV infection and pathology. Although this species is well-known for the susceptibility to HIV that it shares with humans [34, 35], it is not widely appreciated that macaques naturally display variation in susceptibility to a broad spectrum of diseases and disorders that mimic those found in humans, e.g., dyslipidemia, alcohol abuse, macular degeneration, and anxiety [36-42]. While the macaque was identified early as a high priority for assembly of a reference genome, and a draft genome subsequently published in 2007[43], the systematic application of genome-wide data in the macaque to the study of human health and disease has yet to materialize. Since 2007, next-generation sequencing technology has speeded the collection of genomic data at steadily decreasing cost, but only a relatively small number of additional macaque genomes have been explored for variation, and none have yet been systematically applied to the problem of human disease. This is unfortunate, given that rhesus macaque colonies at many of the national primate research centers constitute a powerful resource for genetic analysis of disease that is on par with many human genetic isolates, due to their maintenance in large outbred and pedigreed colonies within a homogeneous environment. Here, in order to catalyze the application of genomic data in macaques to the study of human disease, we present an approach that will make feasible the collection of accurate, dense genome-wide sequence data in large numbers of pedigreed macaques without the need for expensive whole-genome sequence data on all individuals.

Our approach is based on using a low-cost reduced representation sequencing method (genotyping-by-sequencing, GBS), to facilitate pedigree-based imputation of dense marker genotypes from selected individuals with whole genome sequence data. In this study, we evaluated the ability of 2 candidate restriction enzymes (*BglII* and *PstI*) to produce genomic fragments for GBS, using both *in silico* and empirical methods. When compared to *BglII*, we show that *PstI* produces substantially larger numbers of high-quality SNVs that are supported by greater sequence read depth. Further, we found that *PstI* libraries provided sufficient coverage over more than twice the number of high-quality variants needed to generate the sparse “framework” markers required to support imputation. This is important because fluctuations in sequencing coverage among individuals are a known characteristic of the GBS method [32], resulting in the frequent inability to call genotypes at all sites in all individuals. Thus, *PstI* produces far more high-quality SNVs than are actually needed, which increases the likelihood of observing a minimum number of framework markers in every individual. Ultimately, the extent of coverage provided by *PstI* allowed us to impute genotypes at ~14.4 million SNVs over all 20 autosomes, using only 4 individuals with WGS and 12 individuals with GBS, at a median 85-88% accuracy throughout our 16-member pedigree. This approach could be applied at a relatively reasonable cost to other managed or natural colonies of Indian-origin rhesus macaques with pedigree information, and potentially to similar groups of other macaque subspecies.

The selection of individuals for WGS that will maximize the accuracy of imputed genotypes throughout the pedigree is a critical component of this approach. We compared GIGI-Pick [14], a pedigree-based statistical approach to prioritizing subjects for WGS, to two other common heuristic methods for selecting individuals for WGS, including sequencing only the most recent generation of the pedigree (“Bottom of Pedigree”), and sequencing only pedigree founders (“Founders”). Due to the small size of our sample pedigree, we used 3 individuals with WGS to impute genotypes at our streamlined set of dense markers, into 13 remaining individuals based on their framework marker genotypes from GBS data. We show that while median accuracy is only somewhat greater for the GIGI-Pick approach compared to the other two approaches, many more individuals overall displayed higher accuracy using the GIGI-Pick selection method, as reflected in the strong upward shift of the interquartile range. Importantly, rare alleles were imputed with exceptionally high accuracy using the GIGI-Pick selection strategy, suggesting that this strategy offers powerful support for downstream analysis of rare variant effects on complex traits. It is possible that these 3 selection methods may perform differently for alternative pedigree configurations, e.g., in a more shallow pedigree, sequencing founders or the most recent pedigree members may provide information equivalent to the more formal strategy implemented in GIGI-Pick. However, we note that the GIGI-Pick approach results in a clear advantage even in our small pedigree that extends to only 2 generations, but which includes many of the most common relationship types found in NHP colonies. Our results are consistent with those of Cheung et al. [14], in that the GIGI-Pick selection approach substantially outperformed both the “Bottom of Pedigree” and “Founders” (i.e., “PRIMUS” in [14]) approaches in the ability to impute common alleles, although our results indicate more consistent accuracy with the “Founders” approach than with the “Bottom of Pedigree” approach.

Using the WGS individuals ranked in order of priority by the GIGI-Pick algorithm, we examined the gain in accuracy of imputed genotypes throughout the pedigree achieved by the consecutive addition of from 1-9 whole-genome sequences. Our results demonstrate that there are excellent compromises available that balance sequencing costs and the ability to obtain dense and accurate marker data. While the accuracy of imputed genotypes was greatest when using all 9 individuals with WGS, most of this accuracy was achieved using the first 4 WGS individuals, i.e., at 4 WGS individuals, median accuracy is at ~92.4% but increased only another 0.8% with the addition of the remaining 5 genomes. These results suggest that an optimal tradeoff between the animals selected for WGS and GBS exists at the ratio of 1 individual selected for WGS, per 3-5 relatives selected for GBS, a cost savings of ~67-83% over WGS of all 6 individuals. We note that these estimates of accuracy were based on careful selection of an optimal, and thus reduced, set of dense markers that were available on chromosome 19. While this strategy was used deliberately to reduce the total computational time required for this study, in our subsequent imputation of the full set of dense markers across the genome, median accuracy only decreased by 5-8% for all chromosomes.

The increase in overall accuracy observed with additional WGS individuals was not shared uniformly among all individuals in the pedigree. For example, while there was a large increase in accuracy between 3 and 4 individuals with WGS, all of this change is due to increased accuracy in A, a founder (see Fig. 1). In this scenario, this improvement is almost certainly due to the inclusion of WGS data from F, a child of A. We note that D remained an outlier in the distribution of genotype accuracy throughout virtually all imputation analyses based on the ranking of WGS individuals by the GIGI-Pick algorithm. This may be due to the limited initial selection of WGS individuals located in the far right lineage, i.e., only when K, P, and C are added to J and used for imputation does accuracy rise for D. This result is consistent with the GIGI-Pick approach, which balances the selection of closely related individuals within the pedigree to facilitate phasing of genotypes, with the selection of more distant relatives to increase the chance of observing unique founder alleles [14]. Because of this compromise, we note that while sequencing only pedigree founders is not the best strategy for maximizing accuracy, founders may remain unselected using the GIGI-Pick approach when the ability to phase genotypes produces more expected allele calls than does the observation of unique founder alleles, for a fixed number of selected individuals. This result also highlights the importance that prior knowledge of phenotypes plays in selecting individuals for WGS. If traits of interest are known to segregate in a particular lineage within the larger pedigree, it may be advisable to manually assign either founders or a close descendant in that lineage for WGS, if neither individual is selected using a more unbiased approach.

The imputation of dense, genome-wide genotypes with high accuracy will allow the unbiased mapping of genetic variants in the macaque genome to disease traits, using either linkage or association approaches. Both of these approaches are important tools in translational research, and should further advance the understanding of human disease already made possible by research in this species. Large pedigreed colonies of macaques, such as the ~4,500 macaques found at the Oregon National Primate Research Center, provide an almost unequaled resource for translational genetic research, due to their multi-generational pedigree structure and the excess of rare and low-frequency variants expected to segregate within this pedigree. Rare and low-frequency variants are expected to play a significant role in human disease [44-47], and we have shown that we can impute these variants in the macaque genome with high accuracy and at a reasonable cost, using the approach we outline here. Moreover, our findings suggest that this approach can be modified to support specific research goals. For example, it may be beneficial to take advantage of the less accurate but greater amount of information provided by the full set of imputed dense markers during initial discovery of variants either linked to or associated with a disease trait, while fine-mapping or replication of a putative trait locus might employ a reduced, optimal set of dense markers likely to provide greater genotype accuracy over a smaller region of interest.

In this study, we demonstrated that it is feasible to obtain comprehensive genome-wide variation at a fraction of the cost of whole-genome sequencing using the GBS method and pedigree-based imputation, which we describe for the first time in a non-human primate genome. However, imputed genotypes will only be as accurate as the underlying whole-genome sequence data and the reference genome to which it is compared. The macaque genome has undergone multiple revisions; however, it remains in draft form and is less complete than many other common model organisms [17, 43]. There are also extremely limited available data on genetic variants in macaques, and no databases with comprehensive information on known variants or population-level allele frequency information are publicly accessible. Both of these factors present obstacles for accurate whole-genome variant calling in macaques, and will thus reduce accuracy for any genotyping approach. Improvements in both of these areas are urgently needed in order to fully realize the value of the macaque as a genetic model of human disease.

## DECLARATIONS

### 1)

#### List of abbreviations

WGS: whole-genome sequencing
GBS: Genotyping-By-Sequencing
SNV: single-nucleotide variant
ONPRC: Oregon National Primate Research Center
MAF: minor allele frequency
GIGI: Genotype Imputation Given Inheritance
CNV: copy number variant
BWA: Burrows-Wheeler Aligner
VCF: variant call format
GATK: Genome Analyzer ToolKit
MCMC: Markov Chain Monte Carlo
ML: most likely genotype calling method
THR: threshold genotype calling method

### 3) Consent for publication

Not applicable.

### 4) Availability of data and material

The GBS and WGS sequence data files supporting the conclusions of this article are in the process of submission to the NCBI Sequence Read Archive, http://www.ncbi.nlm.nih.gov/sra.

### 5) Competing interests

A. Vinson receives compensation as an external consultant to Novo-Nordisk, USA. No other authors declare competing interests.

### 6) Funding sources

This project was supported by the Office of the Director/Office of Research Infrastructure Programs (OD/ORIP) of the NIH (grant no. OD011092). This funding body played no part in the design of the study, in the collection, analysis, and interpretation of data, or in the writing of this manuscript.

### 7) Authors contributions

All authors made substantial contributions to the conception and design of this study. AV, BB, BF, LC, and RCJ wrote or critically revised the manuscript. BB, AV, JL, KN, LC, BF, MR, and ES analyzed or interpreted data. All authors gave approval for the submission of this manuscript.

## 8) Acknowledgements

We owe particular thanks to Dr. Laura Cox of the Texas Biomedical Research Institute for initial discussions of GBS methods. We also thank Dr. Ellen Wijsman for helpful discussion of key issues around imputation in pedigrees, and Dr. Charles Cheung for support with the GIGI suite of software. We thank the ONPRC Bioinformatics service core for initial processing and analysis of sequence data, and the ONPRC DNA Bank for access to NHP samples used in this project. This project was supported by the Office of the Director/Office of

Research Infrastructure Programs (OD/ORIP) of the NIH (grant no. OD011092).

## 9) Authors’ information

Not applicable.

## 10) Endnotes

Not applicable.

## 13) Tables and captions

Not applicable.

